# Frequency and Phase Correction of GABA-Edited Magnetic Resonance Spectroscopy using Complex-Valued Convolutional Neural Networks

**DOI:** 10.1101/2023.05.13.540263

**Authors:** Hanna Bugler, Rodrigo Pommot Berto, Roberto Souza, Ashley D. Harris

## Abstract

**Purpose:** To determine the significance of complex-valued inputs and complex-valued convolutions compared to real-valued inputs and real-valued convolutions in Convolutional Neural Networks (CNNs) for frequency and phase correction (FPC) of GABA-edited Magnetic Resonance Spectroscopy (MRS) data.

**Methods:** An ablation study was performed to determine the most effective input (real or complex) and convolution type (real or complex) to predict frequency and phase shifts in GABA-edited MEGA-PRESS data using CNNs. The best CNN model was subsequently compared to two recently proposed deep learning (DL) methods for FPC of GABA-edited MRS. All methods were trained using the same experimental setup and evaluated using GABA’s signal-to-noise ratio and linewidth, Choline artifact, and by analyzing the reconstructed final difference spectrum. Statistical significance and effect size were assessed using the Wilcoxon signed rank test and Cohen’s d respectively.

**Results:** The ablation study showed that using complex values for the input represented by real and imaginary channels in our model input tensor, with real (conventional) convolutions was most effective for FPC. For the comparative study, the simulated data test set showed that our CNN model that received complex-valued inputs with real convolutions outperformed models it was compared against with a lower mean absolute error (p<0.05). For the in vivo data test set, our model performed similarly to other DL FPC models.

**Conclusion:** Our results indicate that the optimal CNN configuration for GABA-edited MRS FPC uses a complex-valued input and real-valued convolutions. This model outperformed existing DL models on simulated data and performed similarly on in vivo data.

## 1 INTRODUCTION

Magnetic Resonance Spectroscopy (MRS) is a non-invasive method to measure metabolite concentrations in vivo such as gamma aminobutyric acid (GABA). GABA is the primary inhibitory neurotransmitter and is widely studied for its role in the flow and regulation of information between different brain regions^1^. While many metabolites can be quantified from a conventional MRS acquisition, GABA is of lower concentration and its peaks are “hidden” under the peaks of other metabolites of higher concentration. Spectral editing techniques such as MEGA-PRESS^1^, are commonly used to measure GABA. This method acquires two subspectra: ON and OFF. In the ON subspectra, an editing pulse is applied to modulate the signal of interest through scalar coupling. In the OFF subspectra, the editing pulse is not applied. The OFF subspectra is subtracted from the ON subspectra to remove the signal of non-interest (present in both subspectra) while leaving the signal of interest. In the case of GABA, the 3 ppm peak is overlapped by creatine. The ON editing pulse is placed at 1.9 ppm to refocus the coupling of the GABA signal at 3 ppm while not affecting the 3 ppm creatine peak.

MEGA-PRESS acquisitions typically collect several ON and OFF transients to obtain a sufficient signal-to-noise ratio (SNR) in the final reconstructed spectrum for analysis. However, frequency and phase shifts often due to subject motion and/or scanner instabilities cause subtraction artifacts that deteriorate the quality of the data. These shifts are commonly corrected retrospectively using frequency and phase correction (FPC) techniques^2^. Several approaches have been proposed to accomplish FPC, such as peak alignment^3^. While these are widely accepted and used in MRS, they are dependent on signal integrity and decline in performance at lower SNRs^2,3^.

Deep Learning (DL) models for FPC have been recently proposed in literature and these DL methods have outperformed peak fitting and spectral registration methods.^4,5^ DL is a branch of machine learning that uses deep neural networks to perform both feature extraction and prediction. It has been applied to numerous challenges in imaging^6^, and has seen recent success in MRS specifically to address FPC. Tapper et al.^4^ initially proposed a multilayer perceptron (MLP) model for predicting frequency and phase shifts respectively, which obtained better results compared to traditional methods such as Spectral Registration (SR). Ma et al.^5^ improved upon Tapper’s approach by adding convolutional layers to the model for optimized feature extraction. While both models made advancements in FPC, they operate on the real portion or the magnitude value of spectroscopy data, ignoring the complex-value nature of the data.

In the Magnetic Resonance Imaging (MRI) literature, most Convolutional Neural Networks (CNNs) reconstruction models receive as input the complex-valued MRI data stored as a tensor with two channels – one for the real and the other for the imaginary portions of the complex data^4,5^. These CNN models can employ either complex-valued or real-valued convolutions^6,7^. Though intuitively having a CNN model that receives as input the complex-valued data and uses complex-valued convolutions makes the most sense, surprisingly, the state-of-the-art techniques receive as input the complex-valued data but deploy real-valued convolutions^8,9^. A simplified analogy can be made in this case by stating that these reconstruction models are implicitly learning the complex-valued algebra. While previous research in MRS has described the improvement of FPC^10^ by using complex-valued data, systematic analysis of the implementation of complex values into this model was not performed.

In this work, we investigate whether DL models for FPC can be further improved by leveraging the complex-valued information of spectroscopy data, and complex-valued convolutions in CNN models. The specific objectives of this work were to (1) determine the best CNN configuration in terms of input types (real versus complex) and convolution operations (real versus complex) for FPC and (2) compare the effectiveness of a complex valued DL FPC model against recent DL models^4,5^.

## 2 METHODS

### 2.1 MODEL ARCHITECTURE

#### 2.1.1 Ablation Study

To determine the effectiveness of complex inputs and complex convolutions in comparison to real inputs and real (conventional) convolutions for FPC, we designed four variations of the same model (*cf*., Table 1). The four models for the ablation study are:

**Table 1.** Ablation Study Model Configuration Comparison. Four variations of the same base model were used to compare the optimal implementation of FPC. Models vary by three aspects: their input (real or complex), their convolutions (real or complex) and their number of parameters.

| Model | Input | Convolution | Parameter Count |
| --- | --- | --- | --- |
| Real-valued input<br>conventional convolutions<br>(RR-CCN) | Real-valued | Real-valued | ~ 9 M |
| Complex-valued input<br>conventional convolutions<br>(CR-CCN) | Complex-valued | Real-valued | ~ 9 M |
| Complex-valued input<br>complex convolutions<br>(CC-CCN) | Complex-valued | Complex-valued | ~ 9 M |
| Complex-valued input<br>complex convolutions<br>with twice the filters<br>(CC-CCN-wide) | Complex-valued | Complex-valued | ~ 18 M |

- Real-valued input with real (conventional) convolutions (RR-CNN): The input to the model is a tensor with a single channel, which represents the magnitude value for the frequency correction model and the real portion of the complex number for the phase correction model.
- Complex-valued input with real (conventional) convolutions (CR-CCN): The complex-valued input to the model is represented as a tensor with two channels, one channel represents the real portion while the other channel represents the imaginary portion of complex data. Conventional convolutions are used, so there is no restraint on how the kernel and the channels interact, leaving it to the learning algorithm how to treat the complex characteristic of the input.
- Complex-valued input with complex convolutions (CC-CNN): The input is similar to the previous case, but the model uses complex convolutions^8^, which restrains computations within the convolution to follow complex algebra rules. As the complex convolution kernels have twice the number of parameters compared to the real valued convolution kernels, the number of filters is halved to maintain the same number of parameters as the other models.
- Complex-valued input with complex convolutions model with twice the number of filters (CC-CNN-wide): Identical to the CC-CNN model with two input channels (real and imaginary) and complex convolution implementation, but with twice as many filters as the CC-CCN model.

#### 2.1.2 Base Model

Figure 1A shows the base model used for the four variations of models in the ablation study. It consists of two blocks of two 1D convolutions with kernels of size five followed by a max pooling layer of size two. Then a flattening layer and two more dense layers. The model uses rectified linear unit activation (ReLU) functions for the convolutional and dense layers, except for the final output layer, which uses a linear activation function. For the complex-valued models, the ReLU is applied separately to the real and imaginary portions of the data. For the fully connected layers in the complex-valued models, the real and imaginary values are flattened (intercalated) into a single channel.

**Figure 1.**
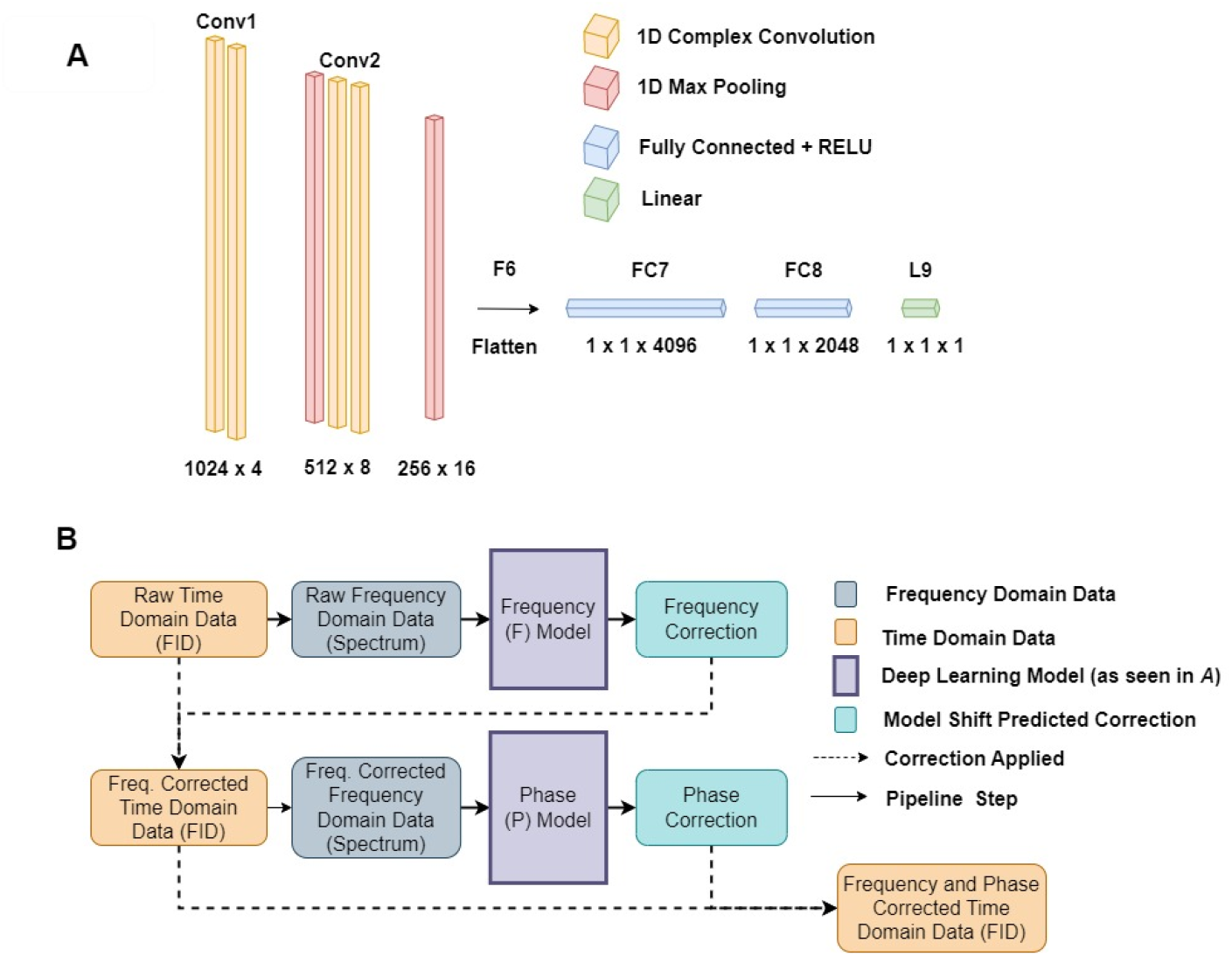
Proposed frequency and phase correction model (CV-CNN). (A) Network architecture is identical for both frequency and phase correction networks. (B) Pipeline for frequency and phase correction. Raw spectral domain data is first passed through the frequency correction network. The frequency shift prediction is then applied inversely to the raw time domain data. The raw spectral domain (frequency corrected) data is subsequently passed through the phase correction network. The phase shift prediction is then applied inversely to the raw time domain data.

#### 2.1.3 Model Architecture and Pipeline

The model pipeline as seen in Figure 1B, equivalent to the one proposed by Tapper et al^4^, consists of two networks connected serially, one for frequency and phase shift prediction respectively, used for both ON and OFF subspectra. A single time domain subspectra (ON or OFF), represented as a free induction decay, is first converted into the frequency domain using the Inverse Fourier Transform and passed into the first network (F-network) used for frequency shift predictions. The frequency offset predicted by the network is then applied inversely to the raw time domain (FID) data to correct for the frequency offset. Following this, the frequency corrected time domain data is converted into the frequency domain using the Inverse Fourier Transform and passed into the second network (P-network) used for phase shift predictions. The phase offset predicted by the network will then be applied inversely to the frequency corrected raw time domain data to correct for the phase offset. Lastly, the time domain data with both frequency and phase correction applied can proceed to subsequent pre-processing steps, such as averaging and subtracting the subspectra, prior to quantification. The complete pipeline has a total of approximately eighteen million trainable parameters as each network, P-network and F-network, contains approximately nine million parameters.

### 2.2 COMPARISON WITH PREVIOUS MODELS

After performing the ablation study and determining the best model among the four proposed variations, the best model was compared to two previously proposed FPC DL models, (1) Tapper et al’s MLP model^4^ and (2) Ma et al’s CNN model^5^. All models were trained under the same conditions and the MLP and CNN model were upscaled to the same parameter count magnitude order for fair comparison by increasing the number of nodes and of filters respectively.

### 2.3 DATA

#### 2.3.1 Simulated Data

Deep learning model performance increases proportionally with the amount of data used for training^11^. As the amount of openly accessible in vivo MEGA-PRESS MRS data is relatively scarce and the true frequency and phase shift values is unavailable for in vivo data, simulated data was used to train and validate the deep learning models.

For the development datasets, 250 GABA+ baseline spectra (ground truth) containing 22 metabolites (alanine, ascorbate, aspartate, beta-hydroxybutyrate, Cr, GABA, glutamine, glutamate, glutathione, glycine, glycerophosphocholine, glucose, myo-inositol, lactate, N-acetylaspartate, N-acetylaspartateglutamate, phosphocholine, phosphocreatine, phosphoethanolamine, scyllo-inositol, serine, taurine)^12^ as well as commonly captured macromolecules (MM09, MM12, MM14, MM17, MM20) and lipid (Lip20) were simulated using the FID-A software^13^. The acquisition parameters of the simulated datasets were: magnetic field strength (3T), MEGA-PRESS variant (FID-A’s MegaPressShaped_fast^13,14^), 14 ms editing pulses at 1.9 ppm and 7.46 ppm with a FWHM= 88.9 Hz, editing interleaving (1 TR), TR/TE = 2 s/68 ms, spectral width (2000 Hz), and the number of spectral data points (N = 2048). The 250 baseline spectra differ in their metabolite concentrations where each metabolite is randomly varied by +/-10% from their averaged reported concentrations in literature^11^.

Each of these basis sets were then used to simulate a set of 320 transients (160 ON and 160 OFF) containing random and linear frequency shifts from −20 to + 20 Hz, random phase shifts from −90 to +90 degrees and random Gaussian amplitude noise. This process was repeated for three different levels of amplitude noise, SNR 10 (good), SNR 5 (fair), and SNR

2.5 (poor). For this work, SNR was defined as the ratio between the amplitude of the creatine peak and twice the estimated standard deviation, using a second order polynomial, between 10 and 11 ppm. Additionally, because the phase models require frequency-corrected data, a version of each dataset without frequency shifts was generated to train the phase models (which will be considered as part of the development dataset). Since previous FPC models tested only on data with a residual water peak, it is hypothesized that these models primarily analyzed the highest peak (water peak) to perform FPC. Therefore, to test the robustness of the models, we repeated the above procedure with simulated data without a residual water peak, resulting in a total of 6 development datasets, as seen in Figure 2, of 250 scans and 320 transients per scan, totalling 80,000 transients per dataset.

**Figure 2.**
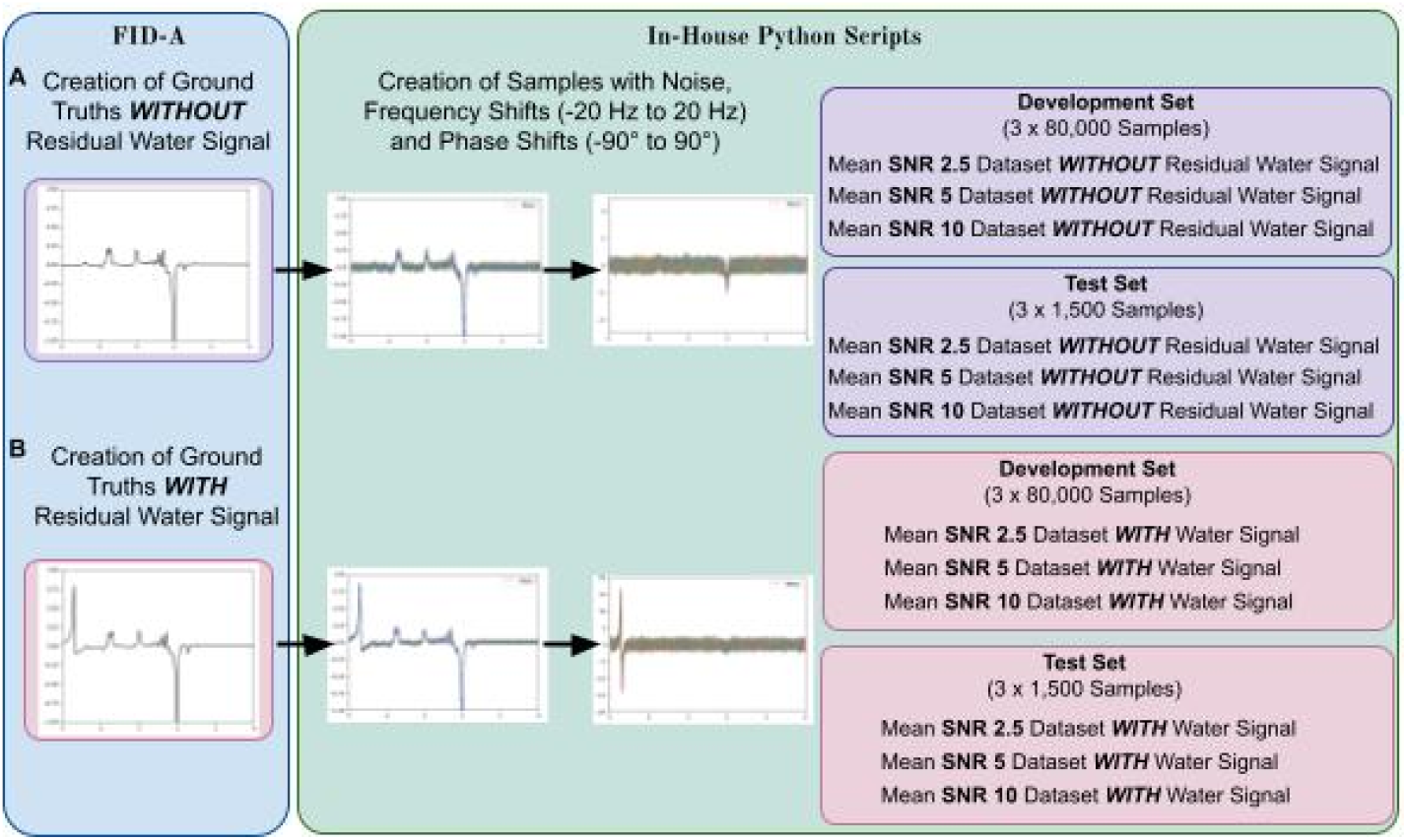
Synthetic (simulated) data generation pipeline for both the ablation and comparative studies. Two sets of baseline spectra containing concentrations of metabolites plus macromolecules and lipids with ten percent variation were generated: (A) one without the presence of a residual water peak and (B) one with the presence of a residual water peak. Ground truths were then loaded by in-house python scripts where each ground truth was used to generate a series of samples (transients) representing an acquisition. Each acquisition (set of transients) was then corrupted to simulate realistic data quality challenges such as adding frequency shifts from −20 Hz to +20 Hz, phase shifts from −90 degrees to +90 degrees to each transient, and Gaussian white noise to generate varying mean SNR of 2.5, 5, and 10.

For the test datasets, similarly to the development dataset, 250 new GABA+ baseline spectra were used to create transients and were subsequently corrupted. From these 250 acquisitions, six transients per acquisition were randomly selected for a total of 1,500 transients. This process was repeated with and without the residual water peak and for each SNR level resulting in six test datasets (two categories of water signal and three categories of SNR) of 1,500 transients.

#### 2.3.2 In vivo Data

Thirty-six in vivo GABA-edited MEGA-PRESS acquisitions from GE scanners of three different sites (G5, G7, G8) with identical parameters from the open access repository Big GABA^15^ were used to compare the three models. Each dataset contained sufficient identical parameters to perform a reliable comparison of the different models.

The acquisition parameters of the in vivo datasets from Big GABA^15^ were: magnetic field strength = 3T, MEGA-PRESS pulse sequence from the ATSM patch, double-echo GRE for B0 shimming, CHESS water suppression, 15 ms editing pulses at 1.9 ppm and 7.46 ppm, interleaving at each TR, TR/TE = 2 s/68 ms, spectral width=2000 Hz, 2048 spectral points, 8-step phase cycle, 320 transients, and 8-channel head coil.

To further evaluate the performance of the models with wider ranges in frequency and phase offsets, artificial frequency and phase shifts were added to the in vivo dataset to create three more datasets, similar to the approach by Tapper et al^4^. The first dataset, defined as having small offsets, had artificial frequency shifts of 0 – 5 Hz and phase shifts of 0-20 degrees. The second, with medium offsets, had artificial frequency shifts of 5 – 10 Hz and phase shifts of 20-40 degrees. The third with large offsets, had artificial frequency shifts between 10-20 Hz and phase shifts of 45-90 degrees.

### 2.4 MODEL TESTING

#### 2.4.1 Simulated Data Training and Testing

All training was performed using the same development dataset and the same 80/20 training/validation division. As there are different datasets which vary by SNR and the presence (or not) of a residual water peak, to assess the impact of model performance for each of these parameters individually, the training and testing was repeated for each dataset. The frequency and phase networks were trained independently, as they have different inputs and outputs, but the data they used was correlated.

The models were trained for 200 epochs using a Mean Absolute Error (MAE). A decaying learning rate that starts at 0.001 and decreases by half every 25 epochs was used. The Adam optimizer^13^ was used and the mean squared error (MSE) was used as the loss function.

After training, test datasets were used to evaluate the prediction pipelines (Figure 2) and obtain frequency and phase correction values, which were compared to the true shift values to calculate the absolute errors. The absolute errors of the frequency and phase predictions for the simulated data were used to calculate the MAE and the standard deviation for each model and dataset. These metrics were used to assess both the hypothesized improvements of using complex-valued convolutions and compare our results with^14,15^. Upon assessing the data, if it follows a normal distribution, a student t-test will be used whereas if it does not follow a normal distribution, a non-parametric test, in this case a Wilcoxon Signed Rank test, will used to determine statistical significance of the absolute prediction error differences between models. Cohen’s d was used to determine the effect size where categorization was defined as small^16^.

#### 2.4.2 In Vivo Data Training and Testing

After the pipelines for all three models were trained on simulated data and tested using the in vivo datasets, the frequency and phase correction values predicted by the model were then applied to the transients. The ON and OFF subspectra were subsequently averaged and subtracted to obtain a GABA difference spectrum.

For qualitative assessment, a sample of the ground truth spectra and the reconstructed spectra by each model using FPC predictions were plotted and compared.

The model’s performance on the in vivo dataset was measured quantitatively using three metrics, Choline (Cho) artifact, GABA SNR and GABA linewidth.

The Q metric proposed by Tapper et al^4^, and then utilized by Ma et al^5^, was used to compare each scan regarding the choline artifact. The Q metric formula is:

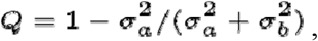

where σ^2^ is the variance of the choline region (3.15 ppm – 3.285 ppm) as a means of quantifying the level of choline subtraction artifact in the final difference spectrum. When comparing spectrum ‘a’ to spectrum ‘b’, Q>0.5 indicates that spectrum ‘a’ obtained a smaller choline subtraction artifact. When Q<0.5, spectrum ‘b’ obtained a smaller choline subtraction artifact.

The GABA SNR was defined as the maximum value in the 2.8 ppm to 3.2 ppm region divided by twice the standard deviation of the 10 ppm to 11 ppm region. Linewidth was defined as the width of the GABA peak where the signal is greater or equal to half of the peak value.

As previously mentioned, if the data follows a normal distribution, a student t-test will be used whereas if the data does not follow a normal distribution, a non-parametric test such as a Wilcoxon Signed Rank test will be used to determine statistical significance of the GABA SNR, GABA linewidth, and Cho artifact differences. Cohen’s d was used to determine the effect size where categorization was defined as small (0.2), medium (0.5) and large (0.8)^16^.

## 3 RESULTS

### 3.1 COMPLEX-VALUED CONVOLUTION ABLATION STUDY

Table 2 shows the MAE results for frequency and phase for the 3 SNR simulations and with and without the residual water peak.

**Table 2.** Ablation Study Results: Mean Absolute Error (MAE) values for frequency and phase predictions on Simulated data with and without a residual water peak for mean SNRs of 2.5, 5, and 10. Results that are bolded indicate the top performing model for the dataset while results that are italicized indicate the second top performing model for the dataset.

| Frequency MAE (Hz) |  |  |  |  |
| --- | --- | --- | --- | --- |
| Dataset | RR-CNN | CR-CNN | CC-CNN | CC-CNN-wide |
| SNR 2.5 | 0.0197 ± 0.0160 | **0.0148 ± 0.0125** | 0.0170 ± 0.0138 | **0.0148 ± 0.0125** |
| SNR 5 | $0.0087 \pm 0.0071$ | $0.0083 \pm 0.0064$ | $0.0082 \pm 0.0072$ | **$0.0076 \pm 0.0064$** |
| SNR 10 | $0.0067 \pm 0.0062$ | **$0.0040 \pm 0.0053$** | $0.0062 \pm 0.0068$ | $0.0048 \pm 0.0053$ |
| No Residual Water Peak SNR 2.5 | $0.5161 \pm 0.4597$ | $0.3211 \pm 0.2930$ | $0.3355 \pm 0.3005$ | **$0.3146 \pm 0.2711$** |
| No Residual Water Peak SNR 5 | $0.1850 \pm 0.1575$ | $0.1400 \pm 0.1233$ | $0.1456 \pm 0.1557$ | **$0.1340 \pm 0.1230$** |
| No Residual Water Peak SNR 10 | $0.0813 \pm 0.0667$ | $0.0612 \pm 0.0489$ | $0.0654 \pm 0.0557$ | **$0.0577 \pm 0.0496$** |
| **Phase MAE (degrees)** |  |  |  |  |
| **Dataset** | **RR-CNN** | **CR-CNN** | **CC-CNN** | **CC-CNN-wide** |
| SNR 2.5 | $0.3174 \pm 0.2735$ | **$0.2374 \pm 0.1913$** | $0.2665 \pm 0.2222$ | $0.2439 \pm 0.2024$ |
| SNR 5 | $0.1409 \pm 0.1170$ | $0.1216 \pm 0.1142$ | $0.1486 \pm 0.1273$ | **$0.1174 \pm 0.0975$** |
| SNR 10 | $0.0802 \pm 0.0705$ | **$0.0681 \pm 0.0558$** | $0.0799 \pm 0.0695$ | $0.0934 \pm 0.0785$ |
| No Residual Water Peak SNR 2.5 | $4.0494 \pm 3.3864$ | $2.6881 \pm 2.2165$ | $2.7854 \pm 2.2475$ | **$2.6738 \pm 2.1474$** |
| No Residual Water Peak SNR 5 | $1.7689 \pm 1.4787$ | $1.2390 \pm 1.0075$ | $1.2865 \pm 1.1061$ | **$1.2367 \pm 1.0267$** |
| No Residual Water Peak SNR 10 | $0.7757 \pm 0.6122$ | **$0.5378 \pm 0.4348$** | $0.6014 \pm 0.4765$ | $0.5841 \pm 0.4670$ |

For frequency and phase shift predictions, each of the four models performed better on datasets with a residual water peak. Similarly, as predicted, all models performed the best on the mean SNR 10 dataset as compared to the lower SNR datasets. Normalization testing revealed that the data followed a non-parametric distribution. Therefore, a Wilcoxon signed rank test was used.

For frequency predictions, the CR-CNN model obtained the best results overall for the datasets with the residual water peak (SNR 2.5: 0.0148 ± 0.0125, SNR 10: 0.0040 ± 0.0053) while the CC-CNN-wide obtained the best results overall for the datasets without the residual water peak (SNR 2.5: 0.3146 ± 0.2711, SNR 5: 0.1340 ± 0.1230 SNR 10: 0.0577 ±0.0496). Each model (CR-CNN, CC-CNN, and CC-CNN-wide) performed significantly (p<0.01) better with a lower MAE for frequency prediction for both the dataset categories (with and without a residual water peak) as compared to the base model (RR-CNN).

Although CC-CNN-wide consistently performed the best or second best overall, due to it having double the amount of parameters as other models, for fair comparison, the base model (RR-CNN) was compared to CR-CNN and CC-CNN models. When comparing the CR-CNN to the CC-CNN model, frequency results were significantly better (p<0.03) with a lower MAE except for both SNR 5 datasets (with and without a residual water peak). When comparing the CR-CNN to the RR-CNN, effect sizes were very small in the SNR 5 with a residual water peak dataset and -small -medium in all others. The effect size between the CR-CNN and CC-CNN-wide models was small to very small in all datasets. Table 3 shows the Cohen’s d effect size values for these two comparisons.

**Table 3.** Ablation Study Effect Sizes: Cohen’s d of the Mean Absolute Error (MAE) for frequency and phase prediction for pairs of most significant comparison.

| Dataset | CR-CNN x RR-CNN |  | CR-CNN x CC-CNN-wide |  |
| --- | --- | --- | --- | --- |
|  | Frequency | Phase | Frequency | Phase |
| SNR 2.5 | 0.245 | 0.244 | 0.011 | 0.036 |
| SNR 5 | 0.059 | 0.138 | 0.039 | 0.012 |
| SNR 10 | 0.120 | 0.120 | 0.031 | 0.207 |
| No Residual Water Peak SNR 2.5 | 0.336 | 0.336 | 0.019 | 0.023 |
| No Residual Water Peak SNR 5 | 0.229 | 0.290 | 0.029 | 0.005 |
| No Residual Water Peak SNR 10 | 0.224 | 0.320 | 0.050 | 0.042 |

Similarly, for phase shift predictions, the CR-CNN model obtained the best overall results for the datasets with a residual water peak (SNR 2.5: 0.2374 ± 0.1913, SNR 10: 0.0681 ± 0.0558) and the CC-CNN-wide obtained the best overall results for the datasets without the residual water peak (SNR 2.5: 2.6738±2.1474, SNR 5: 1.2367 ± 1.0267). For the top two overall best performing models (CR-CNN and CC-CNN-wide) phase prediction MAE for both the dataset categories (with and without a residual water peak) as compared to the base model (RR-CNN), obtained significant (p<0.01) results. When comparing the CR-CNN model to the CC-CNN model, phase results were significant (p<0.03) for all datasets, with the exception of the SNR 5 without a residual water peak dataset. When comparing the CR-CNN to the RR-CNN, effect sizes were small in all datasets. The effect size between the CR-CNN and CC-CNN-wide models was small for the SNR 10 with a residual water peak dataset and very small in the other datasets.

Given that the CC-CNN-wide model has double the amount of parameters in comparison to the other models and thus requires more computational and time resources, and the small difference between the CC-CNN-wide and the CR-CNN, the CR-CNN model was selected as the best performing model and used for the comparison with existing DL FPC models from the literature.

### 3.2 COMPARISON WITH PREVIOUS MODELS

#### 3.2.1 Simulated data testing

Table 4 shows the comparison between the best performing model from the ablation study, the Complex-valued conventional convolution (CR-CNN) model and previous DL FPC models from Tapper et al^4^ and Ma et al^5^ on the simulated datasets.

**Table 4.** Comparative Study Results: Mean Absolute Error (MAE) values for frequency and phase predictions on Simulated data with and without a residual water peak signal for mean SNRs of 2.5, 5, and 10. Results that are bolded indicate the top performing model for the dataset.

| Frequency MAE (Hz) |  |  |  |
| --- | --- | --- | --- |
| Dataset Name | MLP (Tapper et al<sup>4</sup>) | CNN (Ma et al<sup>5</sup>) | CR-CNN |
| SNR 2.5 | 0.0352 ± 0.0446 | 0.0190 ± 0.0154 | **0.0148 ± 0.0125** |
| SNR 5 | 0.0215 ± 0.0279 | 0.0104 ± 0.0095 | **0.0083 ± 0.0064** |
| SNR 10 | 0.0167 ± 0.0204 | 0.0067 ± 0.0059 | **0.0040 ± 0.0053** |
| No Residual Water Peak SNR 2.5 | 0.6495 ± 0.7419 | 0.5696 ± 0.5295 | **0.3211 ± 0.2930** |
| No Residual Water Peak SNR 5 | 0.2421 ± 0.2237 | 0.2359 ± 0.2139 | **0.1400 ± 0.1233** |
| No Residual Water Peak SNR 10 | 0.1113 ± 0.1207 | 0.1031 ± 0.0910 | **0.0612 ± 0.0489** |
| Phase MAE (degrees) |  |  |  |
| Dataset | MLP (Tapper et al<sup>4</sup>) | CNN (Ma et al<sup>5</sup>) | CR-CNN |
| SNR 2.5 | 0.5535 ± 0.6249 | 0.3500 ± 0.3013 | **0.2374 ± 0.1913** |
| SNR 5 | $0.2933 \pm 0.3509$ | $0.1733 \pm 0.1540$ | **$0.1216 \pm 0.1142$** |
| SNR 10 | $0.1546 \pm 0.1699$ | $0.0838 \pm 0.0698$ | **$0.0681 \pm 0.0558$** |
| No Residual Water Peak SNR 2.5 | $5.1314 \pm 4.1875$ | $4.7245 \pm 3.8461$ | **$2.6881 \pm 2.2165$** |
| No Residual Water Peak SNR 5 | $2.2342 \pm 1.8952$ | $2.1183 \pm 1.7688$ | **$1.2390 \pm 1.0075$** |
| No Residual Water Peak SNR 10 | $1.1475 \pm 1.0636$ | $0.9950 \pm 0.8204$ | **$0.5378 \pm 0.4348$** |

Similarly to the ablation study, for frequency and phase shift predictions, each of the FPC models performed better on the dataset with the water signal as compared to the dataset without the residual water peak. In addition, model performance increased with the mean SNR value of the simulated datasets. Normalization testing revealed that the data followed a non-parametric distribution. Therefore, a Wilcoxon signed rank test was used. All results, for both frequency and phase, from the CR-CNN model when compared to others in this comparative study were significant (p<0.01).

For frequency predictions, the CR-CNN model consistently obtained the best results overall for both the dataset with the residual water peak (SNR 2.5: 0.0148 ± 0.0125, SNR 5: 0.0083 ± 0.0064, SNR 10: 0.0040 ± 0.0053) and without the residual water peak (SNR 2.5:0.3212 ± 0.2930, SNR 5: 0.1400 ± 0.1233, SNR 10: 0.0612 ± 0.0489). When comparing the CR-CNN model to Ma et al’s CNN model, which showed the second best performance, the difference in performance was of a medium effect for data containing a residual water peak and a and small to medium for frequency predictions with a residual water peak. Table 5 shows the Cohen’s d effect size values for comparing the CR-CNN model to Ma et al’s CNN model and to Tapper et al’s MLP models.

**Table 5.** Comparison Study Effect Sizes: Cohen’s d of the Mean Absolute Error (MAE) for frequency and phase prediction for pairs of most significant comparison.

| Dataset | CR-CNN x MLP (Tapper *et al*) | CR-CNN x CNN (Ma *et al*) |
| --- | --- | --- |

|  | Frequency | Phase | Frequency | Phase |
| --- | --- | --- | --- | --- |
| SNR 2.5 | 0.221 | 0.316 | 0.478 | 0.534 |
| SNR 5 | 0.171 | 0.276 | 0.487 | 0.504 |
| SNR 10 | 0.131 | 0.157 | 0.457 | 0.463 |
| No Residual Water Peak SNR 2.5 | 0.396 | 0.426 | 0.448 | 0.471 |
| No Residual Water Peak SNR 5 | 0.386 | 0.419 | 0.407 | 0.459 |
| No Residual Water Peak SNR 10 | 0.402 | 0.493 | 0.391 | 0.543 |

For phase shift predictions, the CR-CNN model also obtained the best overall results for both the dataset with the residual water peak (SNR 2.5: 0.2374 ± 0.1913, SNR 5: 0.1216 ± 0.1142, SNR 10: 0.0681 ± 0.0558) and without the residual water peak (SNR 2.5: 2.6881 ±2.2165, SNR 5: 1.2390 ± 1.0075, SNR 10: 0.5378 ± 0.4348) with mean absolute errors larger than those obtained for frequency predictions on similar SNR datasets. Compared to the model by Ma et al that consistently obtained the second best results, the difference in phase prediction errors was of a medium effect size without a residual water peak and small with the residual water peak.

#### 3.2.2 In Vivo Testing

Figure 3A presents results for the GABA linewidth. The datasets with no offsets and small offsets show visually that each model performed similarly while the medium and large offsets dataset showed that our model marginally outperformed the MLP and CNN models. Figure 3B presents results for the GABA SNR. This metric shows the CNN model having the best SNR overall across the different datasets, while MLP performed the worst from no added offsets to medium offsets and the CR-CNN performed the worst on the large offset dataset. However, despite these visual differences, there were no statistically significant differences between models in all cases for all metrics with the exception of the large offset dataset for the GABA SNR metric with very small effect sizes (Cohen’s d<0.2).

**Figure 3.**
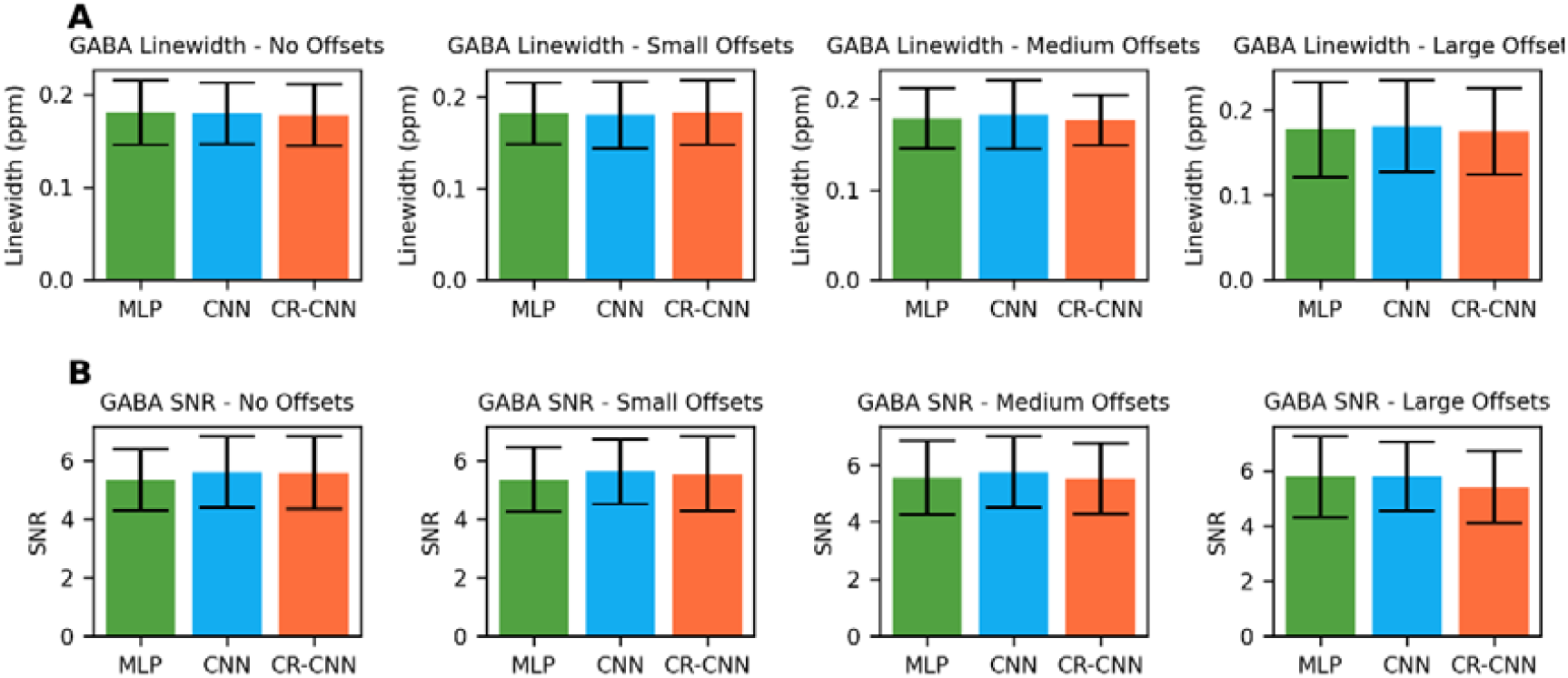
(A) Comparison of model performance by GABA linewidth as calculated by the full width at half max of the GABA peak for in vivo data with (from left to right) no offsets, small offsets, medium offsets, and larger offsets. (B) Comparison of model performance by GABA SNR as calculated by the GABA peak signal over the standard deviation of the noise present between 10 and 11 ppm in vivo data with (from left to right) no offsets, small offsets, medium offsets, and larger offsets.

Figure 4 compares the three models using the Q metric for each scan of the four in vivo datasets. Figure 4A compares the CR-CCN (our) model with Ma et al’s CNN model, Figure 4B compares the CR-CNN (our) model with Tapper et al’s MLP model, and Figure 4C compares the CNN model with the MLP model. We see the average value for Q in all of the graphs is between 0.48 and 0.53 which indicates that there is no model which performs better than the others. This agrees with the graph of Figure 3A, which did not have statistically significant differences.

**Figure 4.**
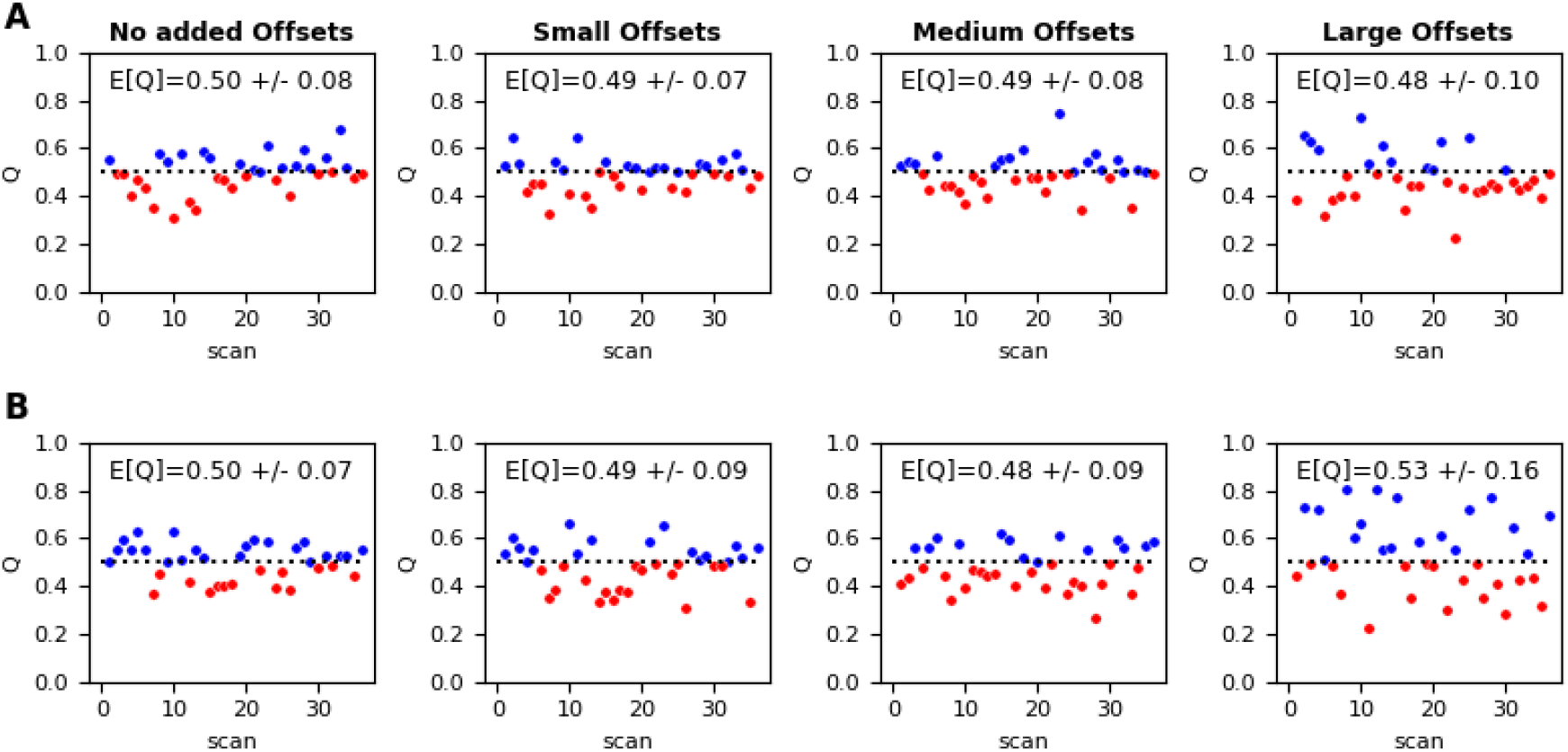
Comparison of model performance for frequency and phase correction using choline artifact Q-plots. Raw in vivo data with different added artificial offsets Q-plot comparison for between the following models:(A) CR-CNN and CNN and (B) CR-CNN and MLP. A blue data point indicates a Q score > 0.5 indicating that the first model obtained a smaller choline artifact as compared with the second model. A red data point indicates a Q score < 0.5 indicating that the second model obtained a smaller choline artifact as compared with the first model. Scores reported on each sub diagram indicate the overall Q score comparing the first model to the second.

In Figure 5, a randomly selected in vivo scan and a randomly selected simulated scan were chosen to evaluate the reconstruction differences between each of the three DL FPC models predicted outputs and the uncorrected raw difference spectrum. The DL FPC models predicted similar frequency shifts. For the GABA peak shape, Ma’s model and our model display more similarity than with the MLP or uncorrected spectra. Analyzing the region just left of the 3 ppm GABA peak, representing the Cho subtraction artifact, we see that they are of similar shape and magnitude in both the CR-CNN and CNN model reconstructions.

**Figure 5.**
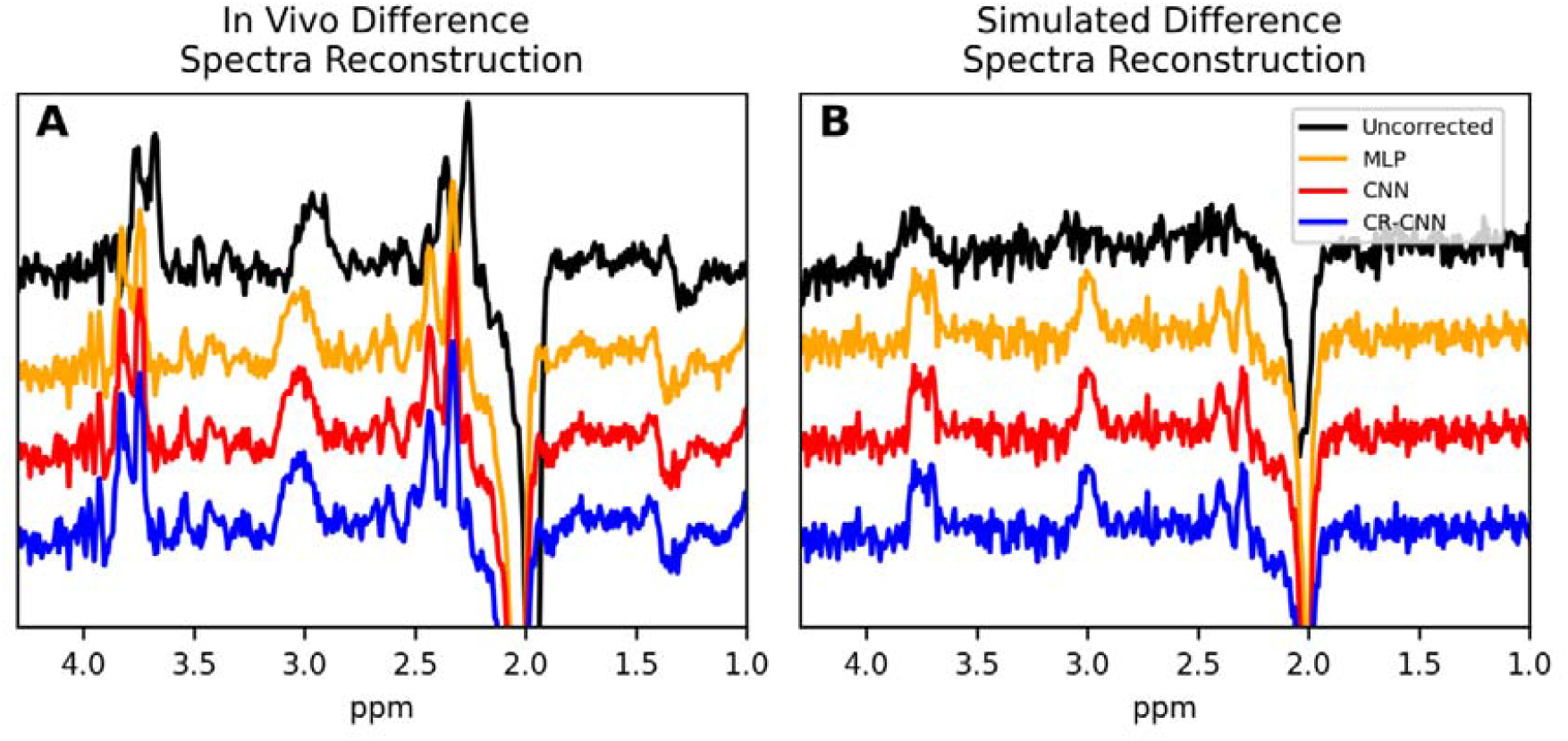
Reconstructed final difference spectra post frequency and phase correction for a single scan of (A) *in vivo* data and (B) simulated data with the residual water peak and SNR of 2.5 using different models for FPC: Tapper et al model (yellow), Ma et al model (red), our model (blue), and uncorrected raw data (black).

## 4 DISCUSSION

Deep learning algorithms for frequency and phase correction of GABA edited-MRS data have been proposed in recent literature to improve data quality and ultimately quantification reliability by reducing noise. Tapper et al initially proposed and tested MLP model to accomplish this task. Ma et al’s built on this work by proposing the CNN model, which improved the accuracy of FPC. We further extend this work by investigating methods to handle the complex values. Our model was consistently more accurate for simulated data and shows comparable performance to the CNN model by Ma et al for the in vivo data using currently accepted and available MRS quality metrics.

### 4.1 ABLATION STUDY

To adequately assess the importance of complex values for FPC, four base models were created to determine whether 1.) the separation of real and imaginary portions of the data by channel and 2.) the introduction of complex kernel operations for convolution would contribute to better feature extraction. The ablation study results demonstrate that separating the input to the model into real and imaginary channels (CR-CNN), instead of using only the absolute value or real part (RR-CNN), improves the FPC results. Changing the convolutions to complex convolutions (CC-CNN), however, did not improve results. The reason for this may be that the CC-CNN models have only half the number of filters, which was necessary in order to maintain the order of magnitude of the parameters with the other models. The CC-CNN-wide model, which keeps the number of filters constant and therefore has double the number of parameters, performed the best in some datasets, but not all.

For this initial study, these results support that while inputting complex-values to the network improves FPC results, using complex convolutions did not result in improvements. This leads us to believe that the feature extraction performed by the convolutional section of the network is better when freely performed in contrast to when constrained to follow complex operation rules. Additionally, as the number of convolutions performed per forward pass is increased by a factor of four when using complex convolutions, the computational effort for the complex convolution models is considerably larger.

### 4.2 COMPARATIVE STUDY

Comparing the CR-CNN (our) model with Tapper et al’s MLP and Ma et al’s CNN model for the simulated datasets, we see that the frequency and phase correction predictions were statistically significantly better for all datasets. This is in agreement with the results from the ablation study, as both MLP and CNN models do not use complex-valued data in their inputs. In contrast to the studies performed by Tapper et al and Ma et al, our study also investigated the role of the residual water peak in FPC results. As predicted, models trained on data with a large positive water peak for FPC outperformed models trained on data without a residual water peak. This is believed to be caused by the deep learning model using primarily the residual water peak to assess frequency and phase changes. Therefore, additional investigations into how deep learning models are learning from the data is required to improve results with superior water suppression.

Comparing the networks for the in vivo datasets and considering the metrics of Cho artifact, GABA SNR and GABA linewidth, we note that there are no significant differences between the DL algorithms, except for GABA SNR in the large offset dataset, in which in which the CR-CNN model obtained a lower SNR, that though significant, is very small in effect size. This lack of differences might be the result of the metrics evaluated, which are the result of 320 frequency and 320 phase predictions and are also affected by other elements of the in vivo data. Thus, the joint precision of 640 predictions are being evaluated together with confounders caused by the metabolite concentrations and noise unrelated to frequency or phase shifts. Additionally, as the differences seen in the simulated data are small in terms of absolute values, they may simply not bring enough effect to alter the metrics evaluated.

When analyzing the reconstructed spectra, we note that each model’s reconstructions are very similar and in agreement with the results of the evaluation metrics.

### 4.3 EVALUATION METRICS

For the purpose of this study, models tested on in vivo data were evaluated using quantitative metrics such as choline standard deviation, GABA linewidth, and GABA SNR and qualitative metrics such as visual inspection of the difference spectrum reconstruction. SNR and linewidth are often used in MRS to assess the quality of spectra and can be indicators of the presence of unwanted subtraction artifacts. In this study, the results interpreted from these metrics in addition to the Q metric did not provide the ability to distinguish quality among model outputs as results from each either lacked agreement and/or were unable to match the precision of the task. Therefore, to further progress in the assessment of machine learning models aimed at improving overall spectrum quality, we believe that better metrics need to be developed. This is however a considerable challenge given the lack of ground truths for in vivo data.

## 5 CONCLUSION

A complex-valued CNN with real and imaginary values contained within two channels is more effective for FPC predictions than utilizing complex convolution. Our CR-CNN model outperformed existing DL FPC models for edited MRS on simulated data and performed equivalently on in vivo data.

